# Soluble guanylyl cyclase subunits act as Hsp90 co-chaperones to ensure the expression and functional maturation of hemeproteins in mammalian cells

**DOI:** 10.64898/2026.08.26.747375

**Authors:** Pranjal Biswas, Yue Dai, Arnab Ghosh, Priya Das Sinha, Dhanya T. Jayaram, Saurav Misra, Dennis J. Stuehr

## Abstract

The cofactor Fe-protoporphyrin IX cofactor (heme) performs many functions in biology. Animal cells must stabilize their newly generated heme-free (apo)-hemeproteins and deliver mitochondrial heme to them so they can mature to functional form. Glyceraldehyde 3-phosphate dehydrogenase (GAPDH) typically accomplishes the heme deliveries, and for many apo-hemeproteins, heat shock protein 90 (Hsp90) drives their heme insertions. We previously observed hemeproteins express poorly in a cell line (COS-7) that does not express soluble guanylyl cyclase (sGC), a heme-binding enzyme that typically functions through its cGMP generation. To understand sGC involvement, we expressed four hemeproteins, Hemoglobin beta (Hbβ), Myoglobin (Mb), Indoleamine 2,3-dioxygenase 1 (IDO1), and Tryptophan 2,3-dioxygenase (TDO) in a cell line expressing sGC (HEK293) or in two cell lines (COS-7, DU145) that do not. We assessed hemeprotein expression levels, their abilities to acquire heme, and when relevant if these facets could be rescued by co-expressing individual sGC subunits, including variants with defects in either sGC heme binding, Hsp90 association, heterodimerization, or cGMP production. We found that co-expression of either sGC subunit was essential for three of the four apo-hemeproteins to accumulate in the COS7 and DU145 cells and acquire heme. This did not involve heme binding, heterodimer formation, or cGMP generation by the sGC subunits, and instead depended on a subunit’s ability to recruit Hsp90 and GAPDH to the apo-hemeproteins via their own Hsp90 binding. Recruiting Hsp90 and GAPDH to apo-hemeprotein clients to ensure they can accumulate and mature to functional form broadens our understanding of sGC and Hsp90 functions in biology.

## Introduction

Many proteins utilize the redox-active cofactor Fe-protoporphyrin IX (heme; hemeproteins) to perform essential biological functions in electron transfer, small molecule transport, catalysis, and signaling (1). Because mitochondria are the source of heme in higher animals and hemeproteins are generated in their heme-free (apo) forms, cells face fundamental challenges to stabilize the apo-hemeproteins and deliver their mitochondrially-generated heme. Our work has uncovered important roles for glyceraldehyde 3-phosphate dehydrogenase (GAPDH) and heat shock protein 90 (Hsp90) in enabling the delivery and insertion of mitochondrial heme into numerous hemeproteins that function in cells outside of the mitochondria (2). In general, the mechanism involves cytosolic GAPDH (3) directly obtaining heme from the mitochondria (4) and safely trafficking it to apo-hemeprotein recipients (2, 5–7), which in most but not all cases (7) are in complex with Hsp90, which is ubiquitously expressed in cells and broadly used to stabilize numerous protein clients and assist in their final folding (8, 9). In the current context, the Hsp90 that is associated with the apo-hemeproteins acts to drive their heme insertions in an ATP-dependent process (10), but whether the Hsp90 association also stabilizes the apo-hemeproteins, and what regulates their association, is still unclear.

During our studies of heme trafficking to apo-hemoglobin-β (Hbβ) and apo-myoglobin (Mb), we found they expressed very poorly in a particular monkey cell line (COS-7) that is reported not express the enzyme soluble guanylyl cyclase (sGC; EC 4.6.1.2), and we found that their expressions could be rescued by sGC co-expression (11). Normally, sGC is widely expressed in all cell and tissue types and regulates diverse biological processes by virtue of its cGMP generation (12–14) (15–17). Structurally, the mature sGC enzyme is a heterodimer comprised of α and β subunits, with the β subunit binding a heme that allows it to bind NO (18), which in turn stimulates cGMP generation (19) (13, 20). Thus, we surmised at the time that the beneficial effect of sGC co-expression was likely related to its cGMP generation, which is known to have positive effects in erythropoiesis (21).

To better understand how the presence of sGC was impacting the hemeprotein expressions, we here performed experiments in which four different hemeproteins (Hbβ, Mb, Indoleamine 2,3-dioxygenase 1 (IDO1), and Tryptophan 2,3-dioxygenase (TDO)) were individually expressed either in a human cell line that naturally expresses sGC (HEK293) (11), or in the COS-7 monkey and DU145 human cell lines, which do not naturally express sGC (22, 23). We measured the abilities of each hemeprotein to be expressed along with their abilities to acquire heme in these different cell types, and if relevant, examined if their expression levels and heme acquisitions could be rescued by co-expressing the individual sGC subunits, including several sGC subunit variants that have specific defects in key sGC properties (heme binding, Hsp90 association, heterodimerization, or catalysis of cGMP production). Our results revealed that co-expression of either sGC subunit were required to stabilize the three “Hsp90-dependent” apo-hemeproteins against proteolysis and to enable their heme incorporations. Remarkably, this did not involve or rely on sGC heme binding, heterodimer formation, or cGMP generation, and instead involved the ability of either sGC subunit to recruit Hsp90 to the apo-hemeproteins, which in turn ensured their stabilization and maturation to functional form. This new non-catalytic role for the sGC subunits is unexpected and stands to broaden our understanding of how sGC and Hsp90 function in hemeprotein biology.

## Results

### sGC co-expression enhances the protein expression and activity of IDO1 but not TDO

We separately expressed FLAG-tagged versions of the hemeproteins IDO1 (IDO1-FLAG) or TDO (TDO-FLAG) in either a human cell line that naturally expresses sGCα or β subunits (HEK293), or in monkey and human cell lines reported to have no sGC expression (COS-7 and DU145, respectively) (11, 22, 23). We verified the presence or absence of sGC expression in the three cell types and confirmed that they all express similar levels of GAPDH and Hsp90 (Fig. 1A-C). The expression level of IDO1-FLAG was much lower in the two sGC-free cell lines compared with its expression in the HEK293 cells, whereas the levels of TDO-FLAG expression were similar in all three cell lines (Fig. 1A-E). To test if a lack of sGC expression in the COS-7 and DU145 cells was related to their low IDO1-FLAG expression levels, we transiently expressed either the α or β subunit of sGC along with either the IDO1-FLAG or TDO-FLAG. The co-expression of either the sGCα or β subunit was successful (Fig. 1A, B) and led to 5-fold increases in their levels of IDO1-FLAG protein expression (Fig. 1A, B, D, E) and to six-fold increases in their IDO1 activities (Fig. 1F, G, H) such that both parameters now equaled the IDO1 expression and activity levels recorded for HEK293 cells. In comparison, co-expressing either sGC subunit in the COS-7 or DU145 cells did not alter their TDO-FLAG protein expression levels, their expression levels of Hsp90 or GAPDH (Fig. 1A, B, D, E), nor did this increase their TDO activities, which remained at levels like the TDO activity seen in the HEK293 cells (Fig. 1F, G, H). Thus, IDO1-FLAG expression in the COS-7 and DU145 cells, but not that of TDO-FLAG, was dependent on cell expression of sGC, and co-expressing either sGC subunit in the COS-7 or DU145 cells was equally effective in rescuing boosting their IDO1-FLAG expression to normal levels seen in the HEK293 cells, along with causing a proportional increase in its enzymatically active form.

**Fig. 1:**
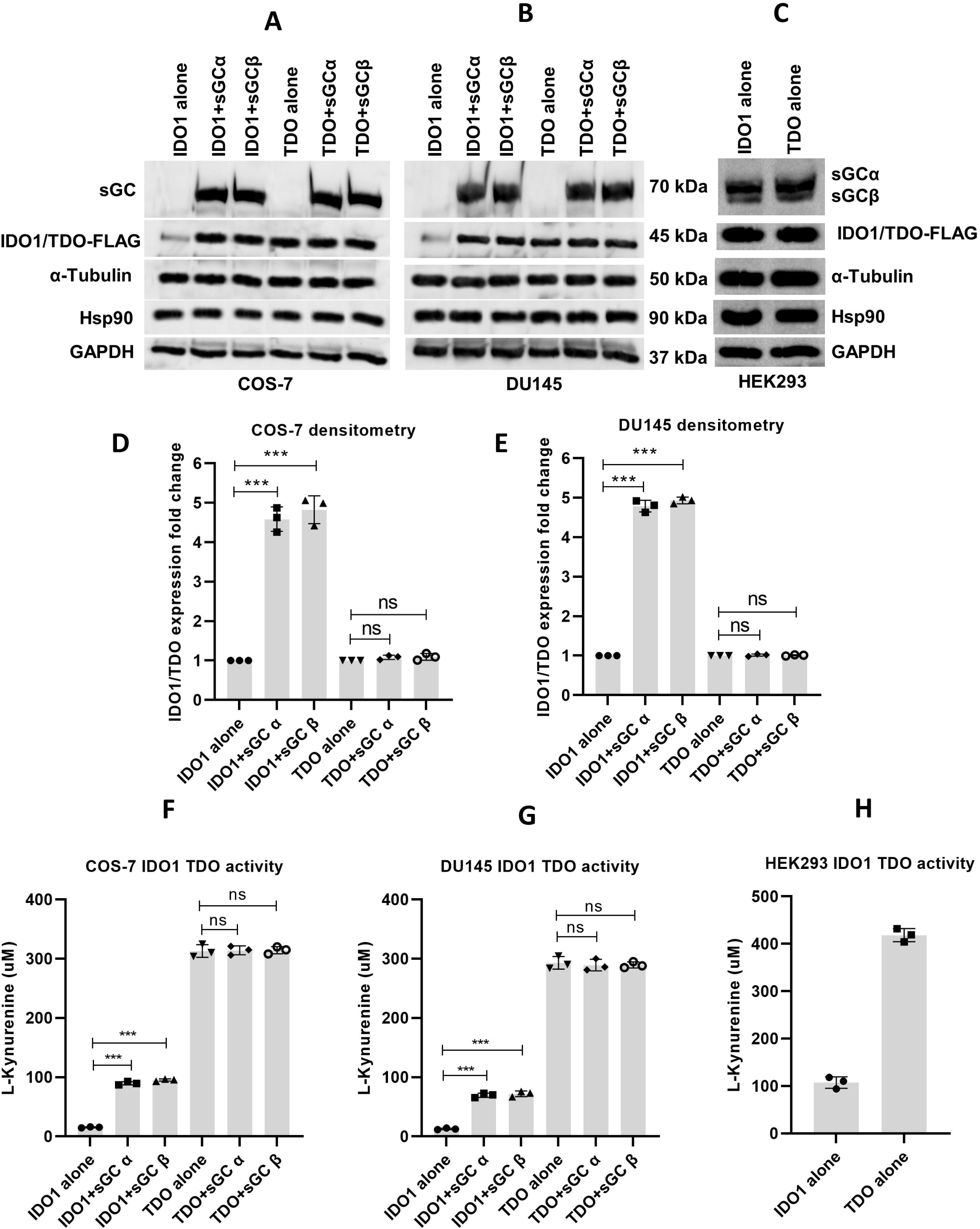
Expression levels and activities of IDO1 and TDO in different cell lines and impact of sGC subunit co-expression. Panels A-C, representative Western blots indicating the protein levels of IDO1-FLAG and TDO-FLAG after they were transiently expressed in COS-7 (A), DU145 (B) and HEK293 (C) cells, along with the cell expression levels of α-tubulin, Hsp90, and GAPDH. Panels D, E, densitometry analysis comparing the relative IDO1 and TDO expression levels in the COS-7 and DU145 cells, respectively, under the indicated conditions of culture. Panels F-H, IDO1 and TDO activities as indicated by the L-Kynurenine accumulation in the COS-7, DU145, or HEK293 cell cultures, respectively, under the indicated conditions. Bars are the mean ± SD, n = 3. ∗∗∗p < 0.001; ns, not significant.

### The positive effect of sGC expression is tied to its Hsp90 binding property

Given these results implicating sGC, we sought to test which if its properties including its heme binding ability, its heterodimerization, its cGMP generation, or its Hsp90 binding ability, were needed to improve IDO1-FLAG protein expression and activity in the two sGC-free cell lines. To do so, we employed several sGCβ and sGCα variants that are defective in one or more of these properties. As listed in Table 1, this included a sGCβ variant that has normal Hsp90 binding but defective heme binding and thus cannot generate cGMP (Y135A R139A; denoted as YARA in our figures) (24), a sGCβ variant that can bind heme and Hsp90 but has defective heterodimerization and thus cannot generate cGMP (Δ204–244+Δ379–408) (11), and a sGCβ variant with defective Hsp90 binding (Δ265-271) that can form a heterodimer but cannot obtain heme in cells, and thus cannot generate cGMP (25). We also generated and tested a sGCα variant (Δ334-340) that was predicted to be defective in Hsp90 binding based on sequence homology between it and the sGCβ subunit. We first confirmed that each sGC variant protein could express well in the COS-7 or DU145 cells and that among them only the sGCβ (Δ265-271) and sGCα (Δ334-340) variants had a significantly reduced association with Hsp90 (Fig. S1A-D). We then individually co-expressed each wild type or variant sGCα and sGCβ subunit with IDO1-FLAG in the COS-7 and DU145 cells via transient transfection and determined after 48 h how the co-expressions impacted the IDO1-FLAG expression level and activity. As shown in Fig. 2A-D, in the absence of either sGC subunit, the IDO1-FLAG showed very weak expression in both the COS-7 and DU145 cells, as we had seen before. Co-expression of the wild type sGC subunits alone or together, or co-expression of most of the sGC subunit variants, led to similar 6-fold increases in the IDO1-FLAG expression level in the COS-7 and DU145 cell lines. The notable exceptions were the Hsp90-binding defective sGCα and β subunit variants, which when co-expressed only supported a 2.5-fold increase in IDO1-FLAG protein expression. The corresponding cell IDO1-FLAG enzymatic activities closely mirrored the protein expression findings: Co-expression of the individual sGC wild type subunits or most of the variants resulted in a 5 to 6-fold boost in activity except for the two Hsp90-binding defective sGC subunit variants, which supported a two-fold boost in activity (Fig. 2E and F). Together, these data indicate that the abilities of either the sGCα and sGCβ subunit to support IDO1-FLAG protein expression and activity in the COS-7 and DU145 cells are linked to their own abilities to bind Hsp90, and do involve or require sGC heme binding, heterodimer formation, or cGMP generation.

**Fig. 2:**
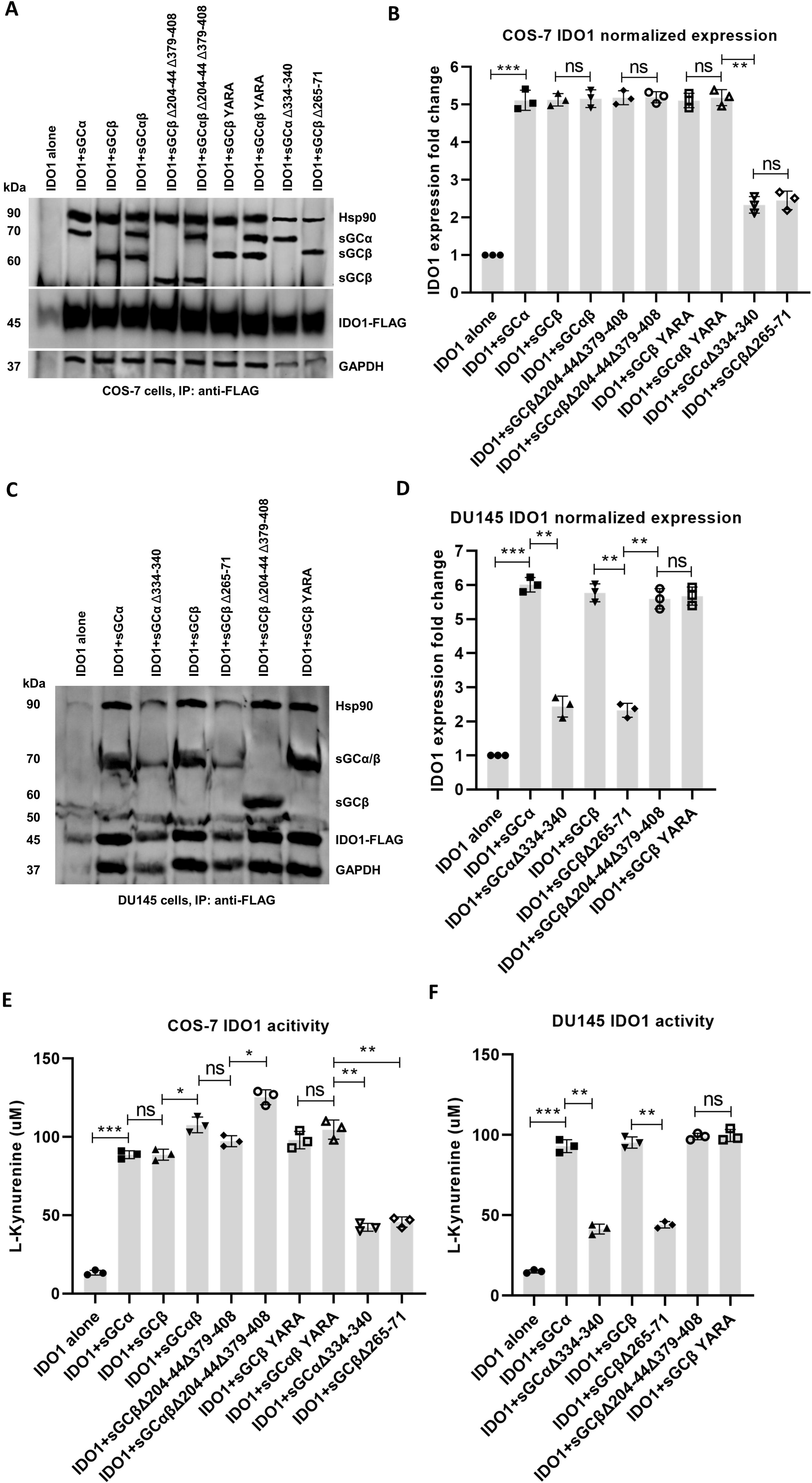
Impact of co-expressing individual sGCα or β subunits or their variants on IDO1 expression level and activity in COS-7 and DU145 cells. IDO1 was expressed in either cell line alone or together with individual sGC α and β subunits or with the indicated sGC subunit variants. Panel A, representative western blot indicating the relative expression levels of the indicated proteins in COS-7 cells. Panel B, normalized protein expression levels based on densitometry analyses of the western blots from A and from two additional independent experiments. Panel C, representative western blot indicating the relative expression levels of the indicated proteins in DU145 cells. Panel D, normalized protein expression levels based on densitometry analyses of western blots from C and from two additional independent experiments. Panels E and F, IDO1 activities as indicated by the L-Kynurenine accumulation in the COS-7 and DU145 cell cultures, respectively, under the indicated conditions. Bars are the mean ± SD, n = 3. ∗∗∗p < 0.001, ∗∗p < 0.01; ∗p < 0.1; ns, not significant.

**Table 1:** List of all mammalian expression plasmids used in the study.

| Plasmid name | Vector | Source |
| --- | --- | --- |
| IDO1-FLAG | pCMV3-C-FLAG | Sino Biological # HG11650-CF |
| TC-IDO1-FLAG | pCMV3-C-FLAG | This study |
| TDO-FLAG | pCMV3-C-FLAG | Sino Biological # HG13215-CF |
| Mb-Myc-FLAG | pCMV6-Entry | Origene # RC213272 |
| TC-Mb-Myc-FLAG | pCMV6-Entry | This study |
| Hbβ-Myc-FLAG | pCMV6-Entry | Origene # RC203258 |
| TC-Hbβ-Myc-FLAG | pCMV6-Entry | This study |
| sGCα1 | pCMV5 | Andreas Papapetropoulos, Greece |
| sGCα1 Δ334-340 | pCMV5 | This study |
| sGCβ1 | pCMV5 | Andreas Papapetropoulos, Greece |
| sGCβ1 Δ265-271 | pCMV5 | Dai et. al., JBC 2019 (25) |
| sGCβ1 Δ204–244 + Δ379–408 | pcDNA3.1 V5-His-TOPO | Andreas Papapetropoulos, Greece |
| sGCβ1 Y135A R139A | pcDNA3.1 V5-His-TOPO | This study |

We next examined if a similar pattern of sGC dependence would hold when Mb-FLAG or Hbβ-FLAG, were expressed in the COS-7 and DU145 cells. As shown in Figs. 3A-D and 4A-D, when expressed on their own the Mb or Hbβ showed weak expression levels in either cell line. Their expression levels were increased by 6 or 7-fold in either cell line when the individual wild-type sGCα and β subunits, the sGCβ subunit defective in heme binding, or the heterodimerization-defective sGCβ subunit were co-expressed, while co-expression with the individual sGCα or β subunits that have defective Hsp90 binding led to lower increases (2 to 4-fold) in Mb or Hbβ expression in the two cell types. Thus, the Hsp90 binding abilities of the sGCα or β subunits were also linked to their enhancing Mb and Hbβ expressions in the COS-7 and DU145 cells, whereas sGC heme binding, heterodimer formation, or cGMP generation did not play a role.

**Fig. 3:**
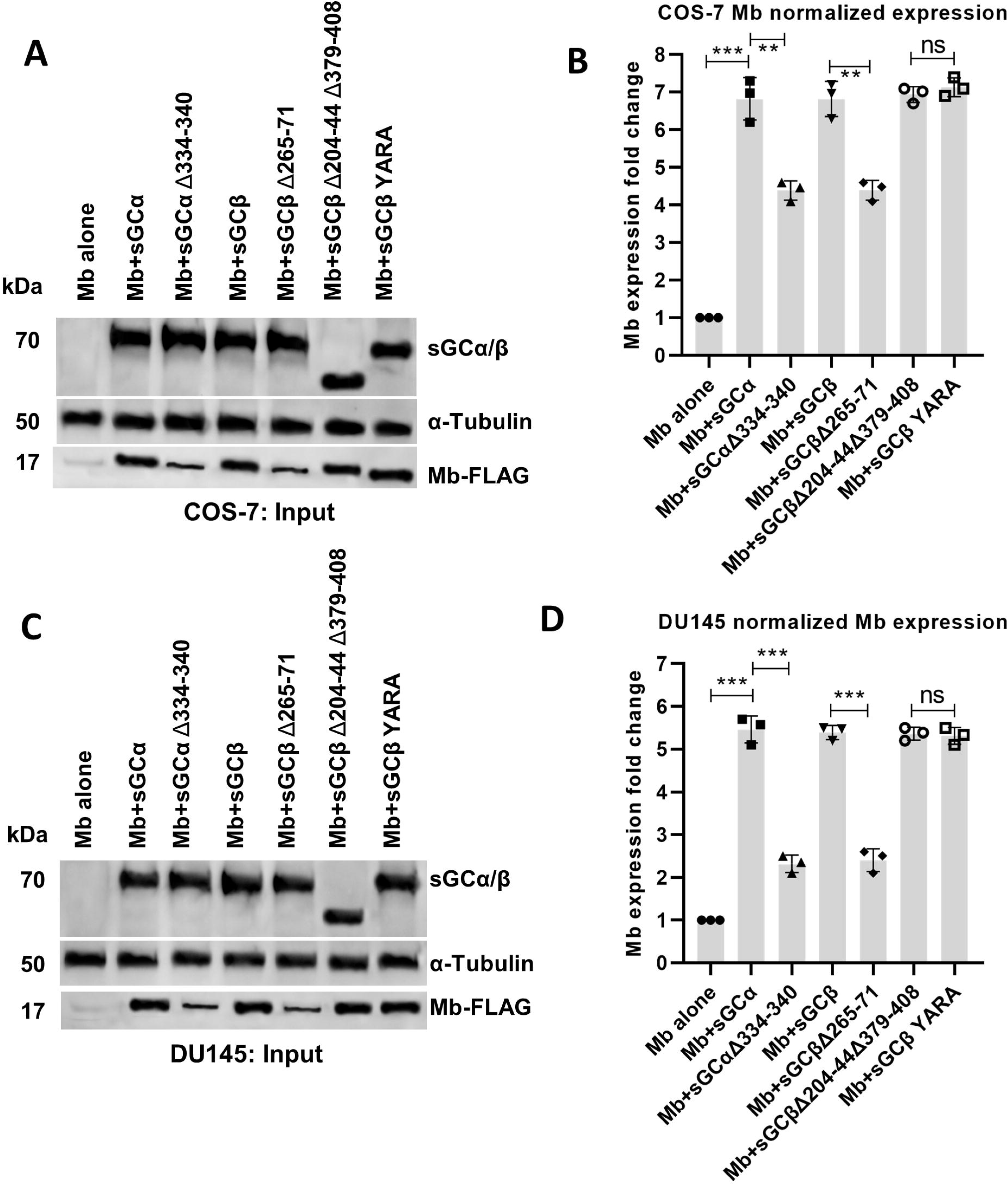
Impact of wild-type or variant sGCα and β subunit co-expression on the expression level of Mb in COS-7 and DU145 cells. Mb was expressed alone or together with individual sGCα and β subunits or their variants in COS-7 or DU145 cells. Panels A and C, representative western blots indicating expression levels of the indicated proteins under the various conditions. Panels B and D, normalized protein expression levels based on densitometry analyses of western blots in A and C and from additional independent experiments. Bars are the mean ± SD, n = 3. ∗∗∗p < 0.001, ∗∗p < 0.01; ns, not significant.

**Fig. 4:**
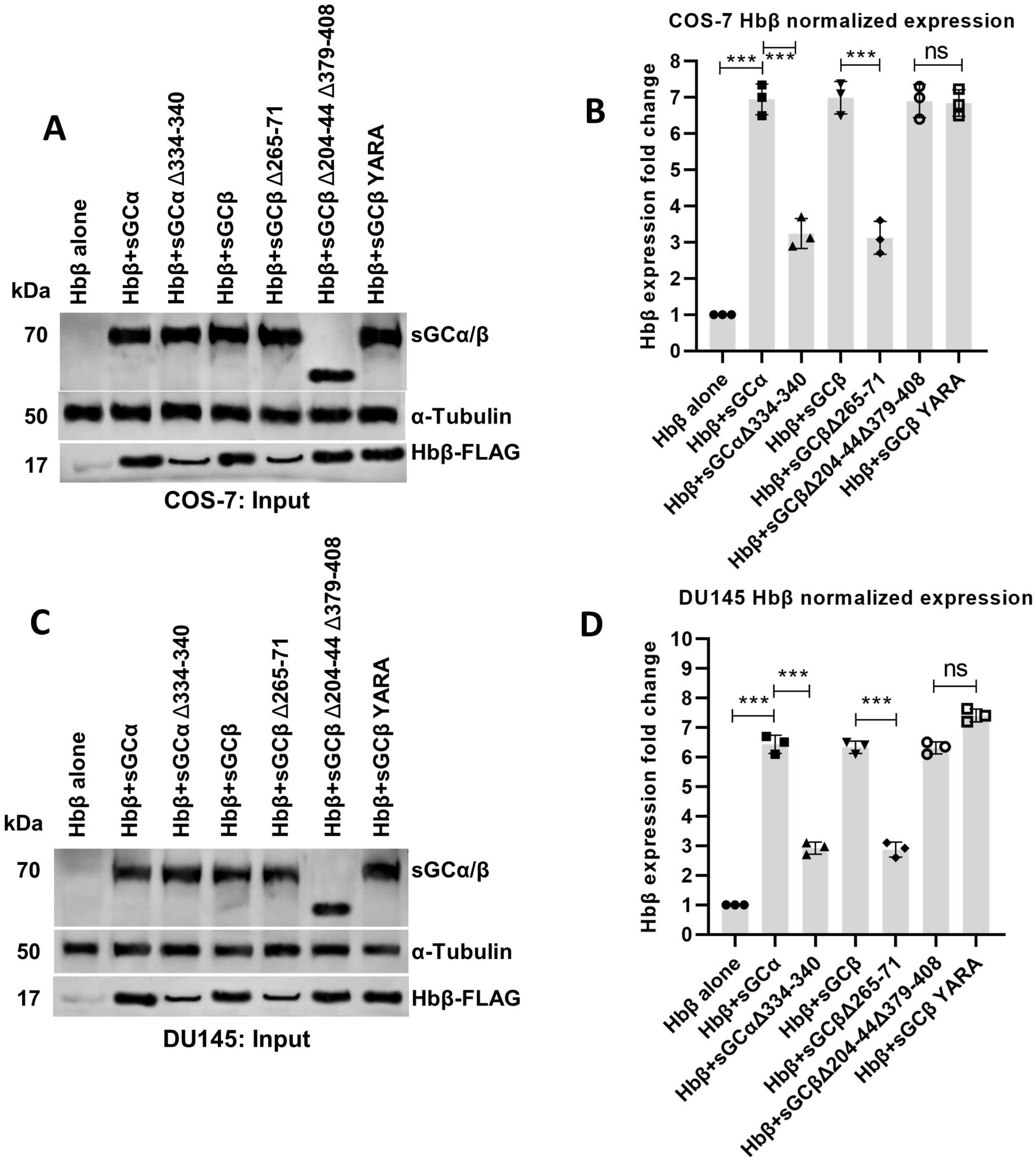
Impact of wild-type or variant sGCα and β subunit co-expression on the expression level of Hbβ in COS-7 and DU145 cells. Hbβ was expressed alone or together with individual sGCα and β subunits or their variants in COS-7 or DU145 cells. Panels A and C, representative western blots indicating expression levels of the indicated proteins under the various conditions. Panels B and D, normalized protein expression levels based on densitometry analyses of western blots in A and C and from additional independent experiments. Bars are the mean ± SD, n = 3. ∗∗∗p < 0.001; ns, not significant.

### sGC acts by enabling Hsp90 association with the apo-hemeproteins

To further probe the mechanism of sGC action, we then attempted to determine how co-expression of the sGC proteins might impact the associations of each FLAG-tagged hemeprotein with Hsp90, GAPDH, and the given sGC protein by performing FLAG IP’s on the various cell supernatants followed by Western blot analyses. Unfortunately, the much lower hemeprotein expression levels achieved in the COS-7 and DU145 cells in the absence of any sGC protein precluded an accurate comparison. Reasoning that the lower hemeprotein “expression” levels seen in these cells in the absence of sGC might actually reflect their poor accumulation due to an increased rate of proteolytic degradation, we tested if inclusion of the proteasome inhibitor bortezomib (26) would alleviate the problem. Including bortezomib at 5 nM during the final 16 h of the hemeprotein transfections maintained good cell viability as judged by trypan blue exclusion (Fig. S2A), and under this circumstance we observed approximate 5-fold gains in the expression levels of FLAG-IDO1, Mb, and Hbβ in the COS-7 cells in the absence of any sGC (Fig. S2B-C). This provided us an approach to examine how either a lack or co-expression of the various sGC subunit proteins might impact each hemeprotein’s associations with Hsp90, GAPDH, and the co-expressed sGC protein.

The panels in Fig. 5A-G indicate the expression levels of the FLAG-tagged IDO1, Mb, and Hbβ proteins when they were individually expressed in the bortezomib treated COS-7 cells on their own or with co-expression of the various indicated sGC subunit proteins, and also compare the relative amounts of Hsp90, GAPDH, and sGC proteins associated with each FLAG-hemeprotein in each circumstance. Representative Western blots are shown in Fig. 5A-C, while the quantifications of normalized band intensities are shown in Fig. 5D-G. The expression level of FLAG-tagged IDO1, Mb, or Hbβ remained similar across all the different culture conditions (Fig. 5D). Without sGC co-expression, little or no Hsp90 or GAPDH was found associated with any of the three hemeproteins. Co-expression with either the wild-type sGCα or β subunit enabled associations between all three hemeproteins with Hsp90, GAPDH, and either sGC subunit, while the sGCα or β variants that have defective Hsp90 binding each supported less associations with Hsp90, GAPDH, and the given sGC variant. Highly similar results were obtained in replica experiments in which DU145 cells were used in place of the COS-7 cells (Fig. 6A-G). Corresponding measures of the IDO1-FLAG activities in the transfected COS-7 and DU145 cells (Fig. 7A and B) showed its activity was directly related to the levels of Hsp90 and GAPDH associated with IDO1-FLAG under the various conditions, consistent with GAPDH and Hsp90 both being essential for apo-IDO1 to obtain heme in cells (7). Thus, expression of either sGCα or β subunit was needed for Hsp90 and GAPDH to associate with each of the three apo-hemeproteins in the COS-7 or DU145 cells. In this process, the sGC subunits also associated with each hemeprotein, and they appeared to function via their own abilities to bind Hsp90.

**Fig. 5:**
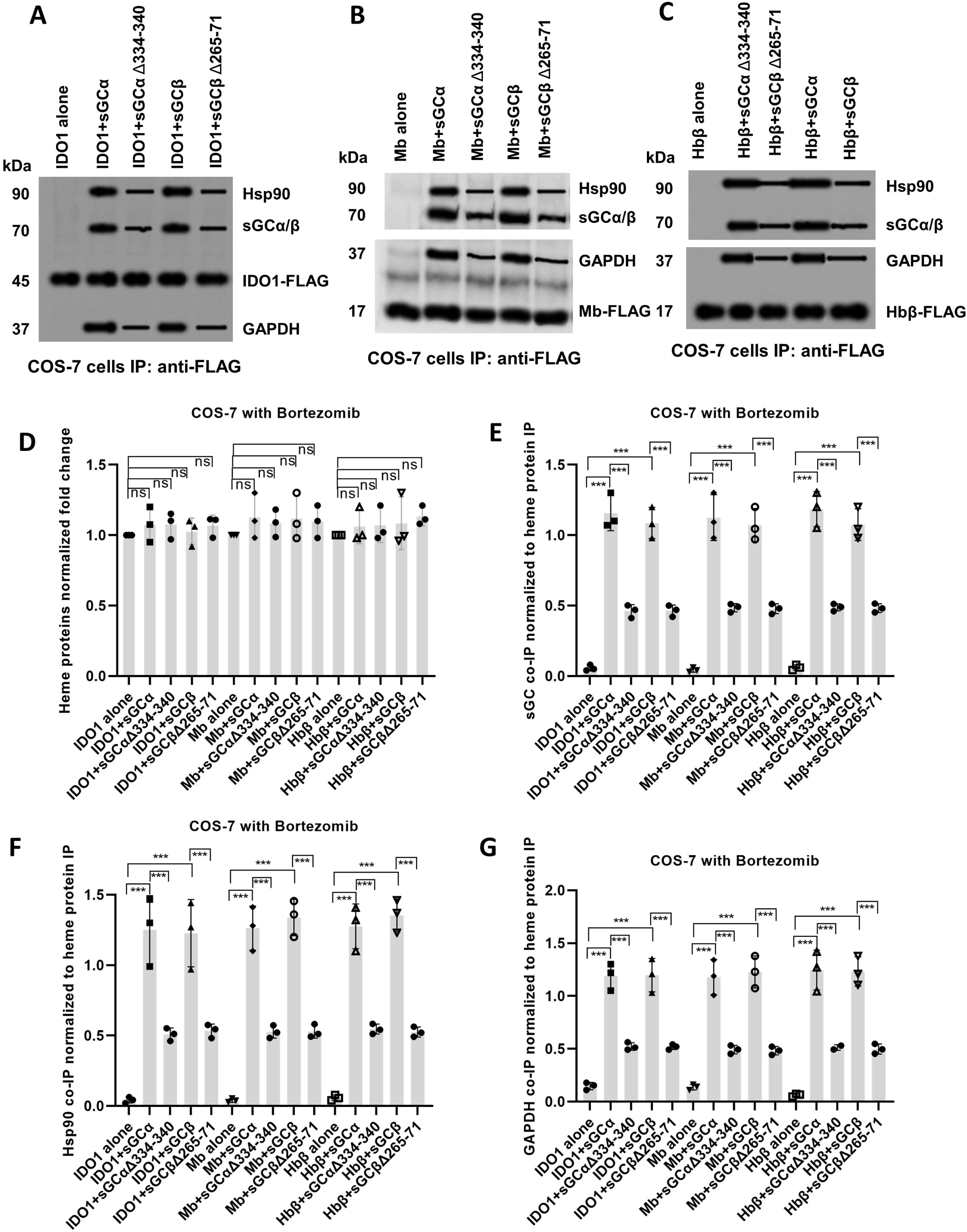
Effect of individual sGC subunit or variant co-expression on sGC subunit, Hsp90, and GAPDH associations with apo-hemeproteins IDO1, Mb, and Hbβ in Bortezomib-treated COS-7 cells. COS-7 cells were treated with 5 nM bortezomib to inhibit the proteasome. IDO1 (A), Mb (B) or Hbβ (C) was expressed alone or with individual sGCα and β subunits or their respective Hsp90-binding mutants. Cell supernatants were then subject to anti-FLAG IP to harvest the hemeproteins and their associated proteins. Panels A-C, representative western blots of the IP samples for each indicated hemeprotein. Panel D, relative heme protein expression levels under each condition, normalized to that for cells expressing the hemeprotein alone. Panels E-G, relative levels of the sGC, Hsp90, and GAPDH proteins associated with each hemeprotein in the IP’s, normalized to the hemeprotein band intensity seen in each sample. Bars are the mean ± SD, n = 3. ∗∗∗p < 0.001; ns, not significant.

**Fig. 6:**
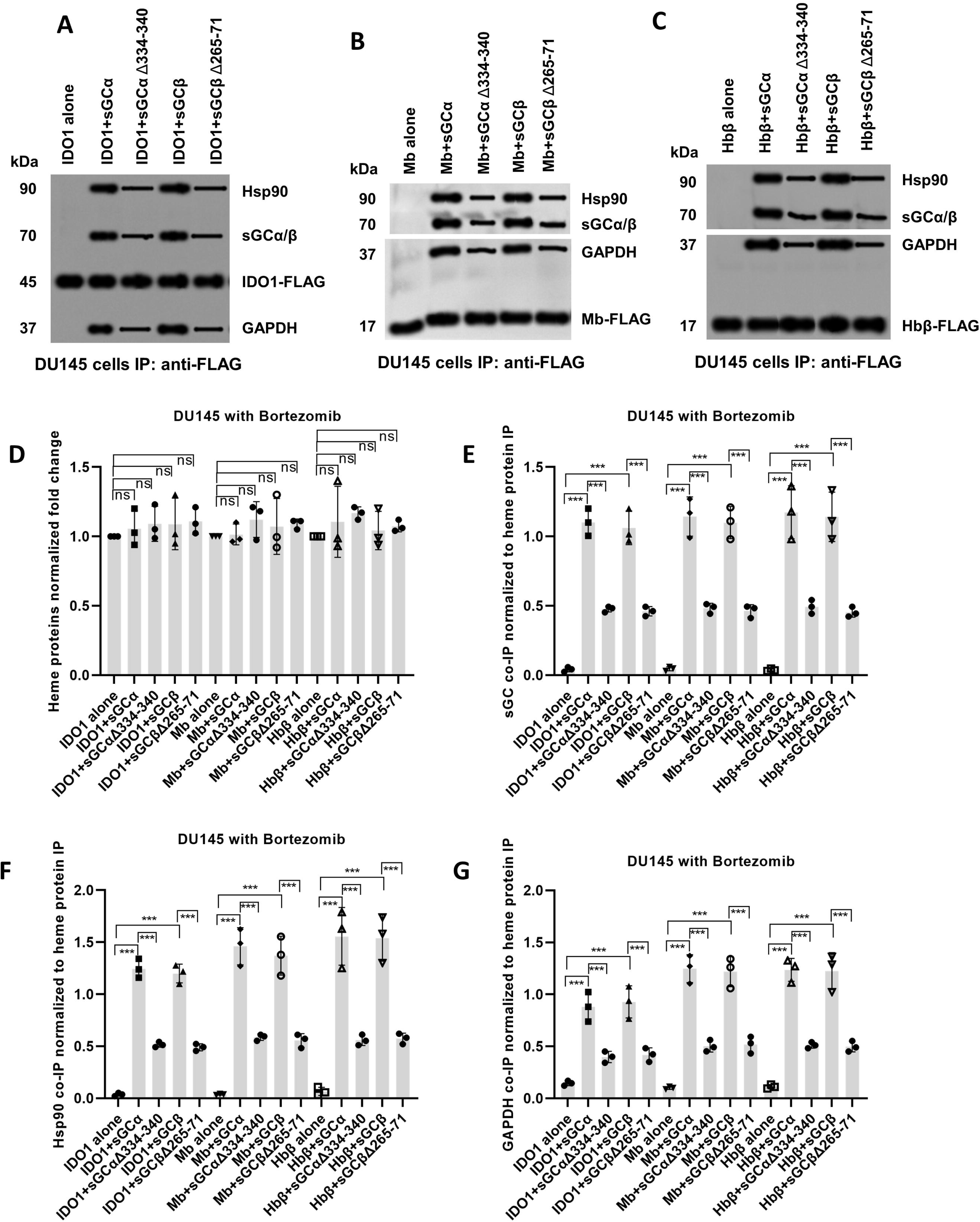
Effect of individual sGC subunit or variant co-expression on sGC subunit, Hsp90, and GAPDH associations with apo-hemeproteins IDO1, Mb, and Hbβ in Bortezomib-treated DU145 cells. The cells were treated with 5 nM bortezomib to inhibit the proteasome. IDO1 (A), Mb (B) or Hbβ (C) was expressed alone or with individual sGCα and β subunits or their respective Hsp90-binding mutants. Cell supernatants were then subject to anti-FLAG IP to harvest the hemeproteins and their associated proteins. Panels A-C, representative western blots of the IP samples for each indicated hemeprotein. Panel D, relative heme protein expression levels under each condition, normalized to that for cells expressing the hemeprotein alone. Panels E-G, relative levels of the sGC, Hsp90, and GAPDH proteins associated with each hemeprotein in the IP’s, normalized to the hemeprotein band intensity seen in each sample. Bars are the mean ± SD, n = 3. ∗∗∗p < 0.001; ns, not significant.

**Fig. 7:**
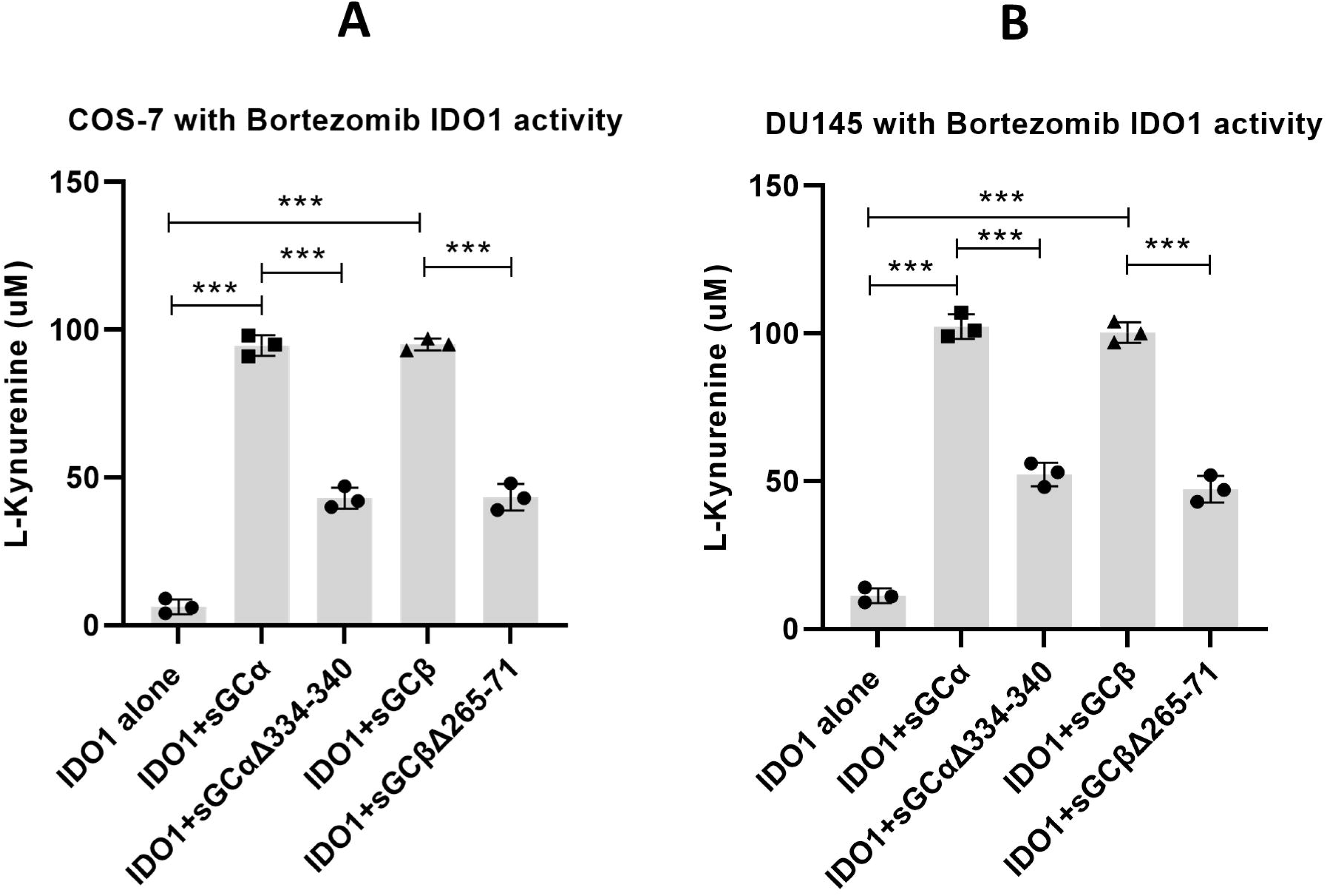
Impact of co-expressing wild type sGC subunits or their Hsp90-binding variants on IDO1 activity in COS-7 and DU145 cells. IDO1 was expressed alone or together with the individual sGCα and β subunits or their Hsp90-binding variants in COS-7 cells (Panel A) or DU145 cells (Panel B). IDO1 activities were assessed by measuring the levels of product L-Kynurenine that accumulated in the cell cultures. Bars are the mean ± SD, n = 3. ∗∗∗p < 0.001.

### sGC enables heme provision into the stabilized apo-hemeproteins

The hemeprotein stabilization that is afforded by bortezomib provided an opportunity to study how sGC co-expression would influence cell heme allocation to apo-IDO1, Mb, and Hbβ, which all depend on GAPDH and Hsp90 for their heme acquisitions (7, 11, 27, 28). To monitor their heme incorporation in live cells, we generated reporter constructs by inserting a tetra-Cys (TC) sequence into each hemeprotein, which is an approach that together with FlAsH-based technology has allowed us to monitor the heme levels in other hemeproteins and in GAPDH in real time in living cells (29, 30). We generated and individually expressed the TC-apo-IDO1, TC-apo-Mb and TC-apo-Hbβ reporter proteins in COS-7 cells that had been made heme-deficient by previous culture with succinyl acetone (SA) (30) and then FlAsH-labeled the TC-apo-hemeproteins. Cell mitochondrial heme biosynthesis was then stimulated by adding the heme precursors δ-aminolevulinic acid and ferric citrate (D-ALA/Fe), and we then followed changes in the cell FlAsH fluorescence emission versus time, where a loss or gain in FlAsH fluorescence intensity indicates a gain or loss in the heme level of the TC-hemeprotein, respectively (30). In the COS-7 cells treated with bortezomib, each expressed TC-apo-hemeprotein gave high FlAsH emission values (Fig. 8A-C), consistent with their accumulating in the cells under this condition. In the bortezomib-treated cells expressing each TC-apo-hemeprotein alone (i.e., without any sGC subunits), little or no FlAsH fluorescence quenching occurred with time after the D-ALA/Fe addition, indicating little or no heme insertion into the TC-apo-hemeproteins took place. In contrast, when these cells co-expressed either the sGCα or sGCβ subunit, we observed fluorescence quenching with time after the D-ALA/Fe addition, indicating that the increase in cell mitochondrial heme synthesis now became coupled to steady heme insertion into the TC-apo-hemeprotein populations (Fig. 8A-C). When we co-expressed the sGCα or sGCβ subunit variants that have an Hsp90 binding defect in place of the wild-type subunits, we observed significantly less fluorescence quenching with time for all three TC-apo-hemeproteins following the D-ALA/Fe addition. Thus, co-expression of either the sGCα or sGCβ subunit was needed for COS-7 cells to allocate mitochondrial heme into TC-apo-IDO1, TC-apo-Mb, and TC-apo-Hbβ, and the capacities of the sGC subunits to do so was directly linked to their own capacities to bind Hsp90.

**Fig. 8:**
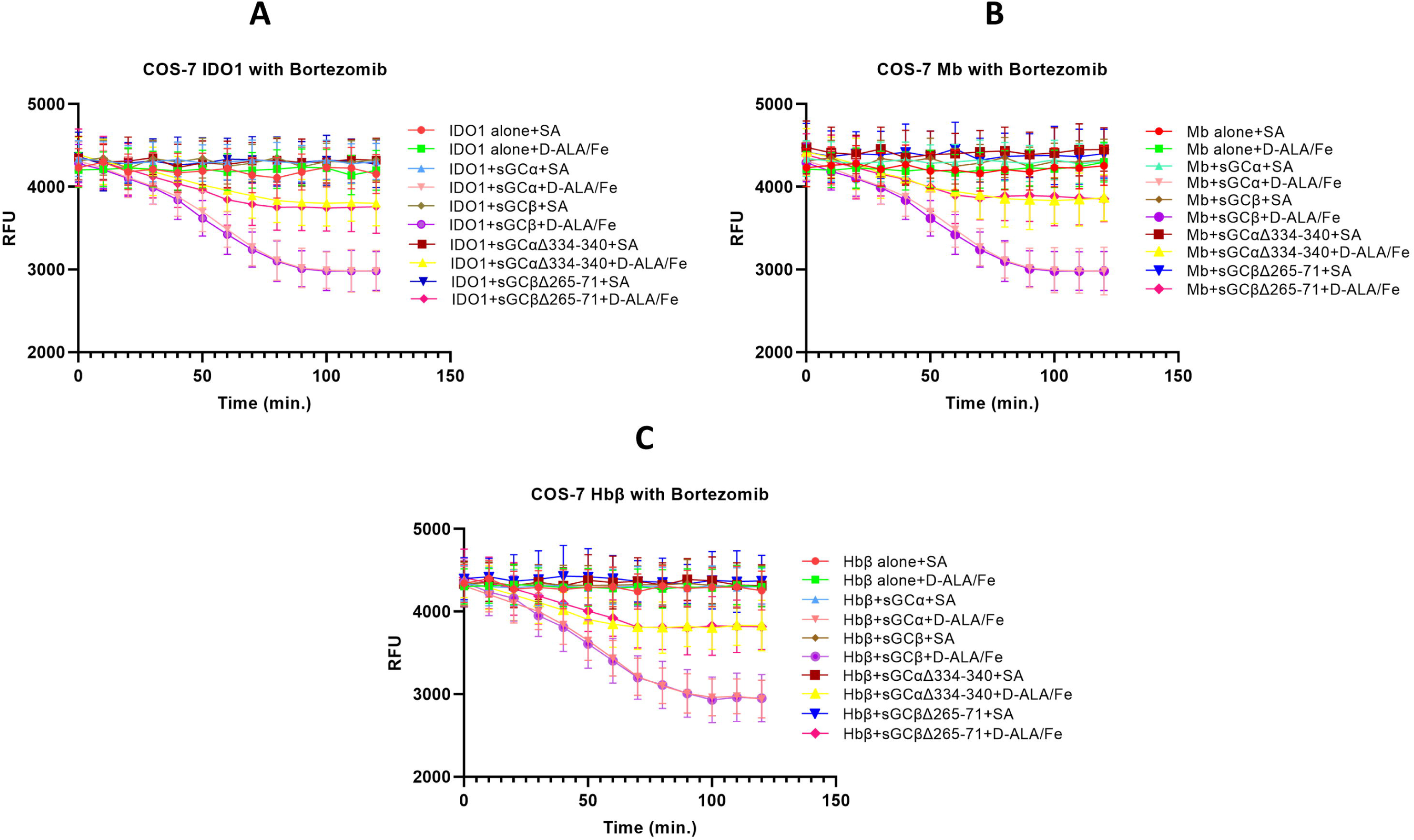
Impact of co-expressing wild type sGC subunits or their Hsp90-binding variants on the abilities of apo-IDO1. COS-7 cells treated with 5 nM bortezomib were transfected to express apo-TC-IDO1, apo-TC-Mb, or apo-TC-Hbβ alone or in combination with wild type sGCα or β subunits and their indicated Hsp90-binding mutants. Following FlAsH labeling of the apo-TC-hemeproteins (in a 96-wellplate), the cells received either buffer alone or buffer plus D-ALA/Fe at time = 0, and the FlAsH fluorescence of each well was recorded vs. time. The extent of heme incorporation into the apo-TC-hemeproteins is indicated by a proportional decrease in the FlAsH fluorescence intensity. Panels A-C, the kinetics of heme incorporation vs. time for each indicated apo-TC-hemeprotein. Bars are the mean ± SD, n = 3.

## Discussion

Our study reveals an unexpected role for sGC subunits in ensuring that certain apo-hemeproteins can accumulate and mature to their functional forms in mammalian cells. Of the four hemeproteins we studied, three (IDO1, Mb, and Hbβ) could barely express in the COS-7 monkey or DU145 human cell lines, which have undetectable sGC expression, unless we pharmacologically blocked cell proteolysis with bortezomib or co-transfected the cells to express either the sGCα or β subunit. In the bortezomib-treated cells, the three hemeproteins could then accumulate normally despite the continued absence of sGC expression, and this allowed us to find that their interactions with Hsp90 and GAPDH and heme acquisitions were prevented unless either sGC subunit was co-expressed. Remarkably, the ability of the sGC subunits to act in this way did not involve or require any core properties of sGC that it normally needs to function in biology, namely heme binding by its sGCβ subunit, sGC heterodimer formation, or cGMP generation. Rather, our findings suggest that the individual sGC subunits enabled Hsp90 and GAPDH to associate with the apo-hemeproteins in the cells. Viewed in this light, the sGC subunits fulfill an important recruitment function that allows these apo-hemeproteins to associate with key cell proteins that both protect them from proteolysis and enable their heme procurement so they can mature to their functional forms in cells. Although this non-enzymatic function for sGC is unconventional, it is in line with sGC co-associating with Hsp90 and the hemeprotein endothelial NO synthase (31) and with sGC participating in Cys-nitroso group transfer between proteins (32).

Mechanistically, the sGC subunits likely stabilize the apo-IDO1, Mb, and Hbβ proteins through their helping to recruit Hsp90. Consider that in their apo forms all three of these hemeproteins bind Hsp90 and require it’s ATPase activity for their heme acquisitions (7, 11, 28). Conversely, the one other hemeprotein we studied (TDO) is distinguished by not binding or requiring Hsp90 for its heme acquisition (7); and unlike the other three hemeproteins, we found that TDO expressed in the COS-7 and DU145 cell lines in the absence of sGC it accumulated to a normal level and matured to its heme-containing, catalytically functional form as well as in the HEK293 cells, which naturally express sGC. These results with TDO ensured that the COS-7 and DU145 cell lines had no inherent defect in supporting hemeprotein expression or maturation and provided the first indication of a possible connection between Hsp90 function and sGC. For the apo-IDO1, Mb, and Hbβ hemeproteins, the importance of sGC-dependent Hsp90 recruitment was further strengthened by our finding that co-expressing sGCα and β variants with documented poor Hsp90 binding affinities (25) supported a proportionally diminished buildup of apo-IDO1, Mb, and Hbβ proteins in the COS-7 or DU145 cells. That Hsp90 recruitment is needed to stabilize the apo-IDO1, Mb, and Hbβ proteins fits squarely with its well-known ability to stabilize many of its client proteins against proteolytic degradation, including kinases, ErbB2, BRaf-V600E, FGFR-G719S, BCR-ABL, and EML4-ALK (33). Hsp90 may also stabilize the sGC subunits themselves, based on a report that the subunits disappear from cells at a faster rate when the ATPase activity of Hsp90 is pharmacologically inhibited (34). The strong influence of sGC subunits on Hsp90 associations is even more remarkable, given that apo-IDO1, Mb, or Hbβ all display robust Hsp90 association when they are expressed in cells or tissues that normally contain sGC. Importantly, ablating or pharmacologically inhibiting the ATP-ase activity of Hsp90, on its own, does not cause it to dissociate from these three apo-hemeprotein clients, which instead takes place after they have incorporated heme (7, 11, 28) (10). This behavior has actually masked the important function of Hsp90 in stabilizing these apo-hemeproteins, which could become apparent only after we performed experiments in sGC-free cells.

At a molecular level, our findings using the sGC subunit Hsp90-binding variants revealed that the ability to recruit Hsp90 to each of the three apo-hemeproteins is heavily dependent on the sGC subunit itself binding Hsp90. Our previous studies showed that major Hsp90 binding elements in the apo-sGCβ subunit are in its Per-Arnt-Sim domain (35). Our current findings suggest that similar Hsp90 binding determinants are also in the sGCα subunit, which had not previously been shown to engage in consequential Hsp90 binding. In the apo-sGCβ subunit, these structural elements enable it to directly bind Hsp90, which in turn is essential for the apo-sGCβ subunit to incorporate heme provided by GAPDH in living cells (36). Thus, our findings suggest that sGC subunit recruitment of Hsp90 to apo-IDO1, Mb, and Hbβ in cells manifests by Hsp90 binding to the same subunit structural elements that need to bind Hsp90 during the process of sGC maturation.

Our IP studies with the FLAG-tagged apo-IDO1, Mb, and Hbβ proteins shed additional light regarding GAPDH recruitment. Consider that in the COS-7 or DU145 cells expressing either sGC subunit, each of the three apo-hemeproteins was associated with GAPDH along with Hsp90 and the sGC subunit. In contrast, in the absence of sGC subunit co-expression, there was no detectable GAPDH association with these apo-hemeproteins in the bortezomib-treated COS-7 and DU145 cells. This was surprising, because in purified systems GAPDH binds well with these apo-hemeproteins, and it is found associated with all three when they are expressed in tissues or cell lines that naturally contain sGC (7, 27). Thus, in addition to enabling their Hsp90 associations, the sGC subunits are needed to recruit GAPDH to the apo-hemeproteins. The GAPDH association is not important for stabilizing the apo-hemeproteins, which build up to normal levels when they are expressed in cells whose GAPDH level has been knocked down (27). This may be because they remain associated with Hsp90-sGC. In a recent investigation of apo-IDO1, we identified GAPDH-apo-IDO1 interface residues, and showed that specific interface residues enable their direct interaction in living cells, which was in turn essential for enabling GAPDH-heme to transfer into apo-IDO1 in the cells (37). This suggests that GAPDH is likely to make direct contact with the apo-hemeprotein when it is bound to Hsp90-sGC.

A working model for sGC participation in hemeprotein maturation that is consistent with the current information is illustrated in Fig. 9. sGC subunit binding to Hsp90 enables Hsp90 to associate with the apo-hemeprotein. At minimum, this creates a 3-protein complex that protects the apo-hemeprotein from proteolysis and may modify its structure. GAPDH is then able to associate to form a 4-protein complex, and this likely allows GAPDH to make direct contact with the associated apo-hemeprotein, thus enabling the GAPDH to perform heme transfer when it contains heme. Once heme is incorporated into the hemeprotein client, the complex dissociates, releasing the mature hemeprotein. Thus, the sGC subunits enable recruitment of, at minimum, two other proteins (Hsp90 and GAPDH) that along with itself form a complex with the apo-hemeprotein to protect it against proteolysis and provide an opportunity for it to receive and incorporate heme to mature to its functional form. This model is likely to be minimalist, because Hsp90 typically interacts with several co-chaperone and heat shock proteins that can influence its interactions and influence on its client proteins to govern their ultimate biological functions (8). Indeed, although the hemeprotein neuronal NO synthase was not studied here, it is known that it’s heme procurement is Hsp90-dependent and influenced by Hsp70 and the co-chaperones Hsp40, Hop, and p23 (38), and we previously found that apo-Mb associates with Hsp70 and the Hsp90 co-chaperones Aha1, Aarsd1, STIP1, and Cdc37 in cells during its heme procurement (11). Based on our current findings, we surmise that the sGC subunits act as additional Hsp90 co-chaperones to ensure Hsp90 and GAPDH can interact with apo-hemeprotein clients so they can safely accumulate in cells and obtain heme.

**Fig. 9:**
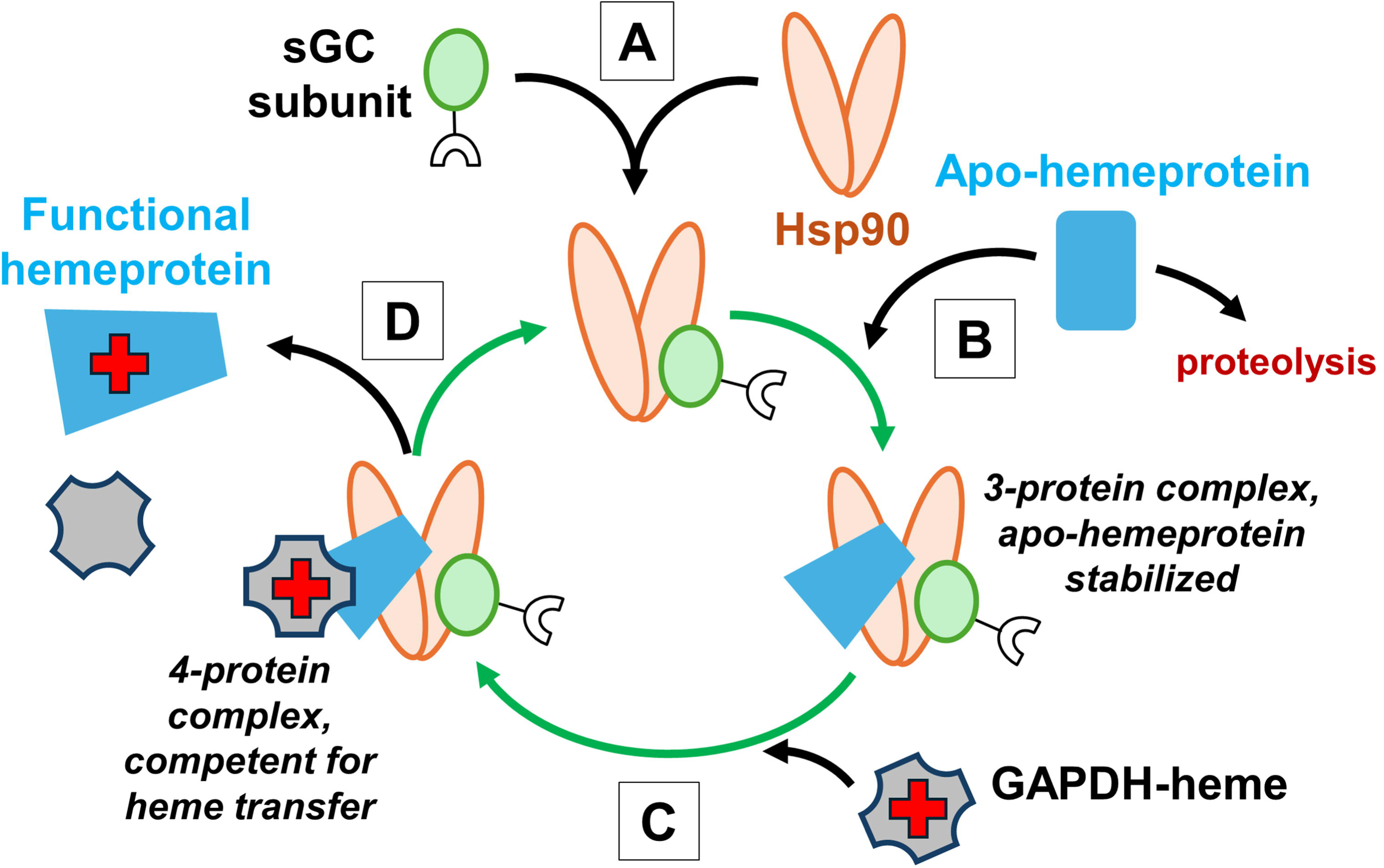
Model for sGC subunit participation in the stepwise maturation of Hsp90-dependent hemeproteins in mammalian cells. Hsp90 binding of a sGC subunit (A) enables it to associate with an apo-hemeprotein (B) to form, at minimum, a 3-protein complex that protects the apo-hemeprotein against proteolysis. GAPDH can then associate (C) to generate, at minimum, a 4-protein complex that can support heme transfer from the GAPDH into the apo-hemeprotein. After the heme transfer is complete the complex dissociates, releasing the functional heme-containing hemeprotein (D) and possibly allowing the Hsp90-sGC species to recycle (green arrows).

In a whole animal context, several studies have reported that sGC can positively influence the expression or function of Hb and certain other hemeproteins (39–41). But in these cases, cGMP generation was either found or presumed to be the primary means of sGC action, which requires that both the sGCα or β subunits be simultaneously expressed. In sGC knockout (KO) studies done with mice, it has only been possible to work with animals missing a single sGC subunit, due to the double KO causing embryonic lethality (42). In such mice with single sGC subunit KO, their sGC-derived cGMP production is severely compromised as expected while the expression of the one remaining sGC subunit remains. In our case, there was a clear redundancy among the two sGC subunits in their abilities to stabilize the apo-hemeproteins. This predicts that such single KO mice would still have normal or near normal buildup and maturation of their Hsp90-dependent hemeproteins, consistent with successful Hb production being observed in the sGCα subunit knockout mice with relatively mild changes in their hematological profiles (40), and also with various outcomes using sGCβ subunit KO mice (43). Perhaps the redundancy displayed by the sGC subunits in stabilizing Hsp90-dependent apo-hemeproteins and in enabling their maturation reflects how critical these processes are for development and viability in higher animals compared to sGC-generated cGMP signaling, which appears to be to be far less critical based on outcomes from the single sGC subunit KO mouse studies. Thus, it follows that a combined lack of sGCα and β expression cannot be tolerated in higher animals due to the deleterious impact this would have on the accumulation and maturation of essential hemeproteins like Hb or Mb. Intriguingly, primary airway smooth muscle or endothelial cells isolated from human donors (44) display a remarkable heterogeneity in their expression levels of the individual sGC subunits, with cells derived from disease cohorts often exhibiting no detectable expression of one or the other sGC subunit, most typically sGCβ. In these cases, besides a compromised cGMP production, the stability and maturation of Hsp90-dependent hemeproteins would become more sensitive to single nucleotide polymorphisms in the remaining sGC subunit. Whether such circumstances exist or track with disease deserves further investigation.

There are several questions to pursue based on the sGC subunits playing key roles in recruiting Hsp90 and GAPDH to apo-hemeproteins to ensure their stability and maturation. For example, what are the molecular details of the Hsp90, sGC subunit, apo-hemeprotein, and GAPDH interactions within the complexes? Why can’t the apo-hemeproteins associate with Hsp90 and GAPDH in cells when sGC subunits are absent, given that they do so readily in their purified forms? What other Hsp90 co-chaperones are involved? How can Hsp90-independent hemeproteins like apo-TDO accumulate in cells and still obtain heme from GAPDH? And finally, is sGC-governance of Hsp90 function restricted to its apo-hemeprotein clients, or is it more broadly involved in governing Hsp90 behaviors toward its numerous other clients? Work can now commence to address these and other questions.

## Materials and methods

### Cell culture

COS-7 cells (ATCC # CRL-1651), DU145 cells (ATCC # HTB-81) and HEK293 cells (ATCC # CRL-1573) were grown in phenol red free DMEM high glucose medium with 10% FBS. Both COS-7 and DU145 cell lines are reported to lack endogenous sGC protein expression (22, 23) and were confirmed by western blot before use in our experiments. Notably, the COS-7 line we originally procured for this study was found to express detectable levels of the sGC subunits, and this confounded our initial experiments. This variability may help explain why IDO1 was previously reported to have normal expression and activity in COS-7 cells (45, 46).

### Plasmids and transfection

Mammalian expression plasmids for rat sGC (sGC α1 WT, β1 WT, and sGC β1 Δ204–244 + Δ379–408 mutant were gifts from Dr. Andreas Papapetropoulos (University of Athens, Athens, Greece). sGC α1 Δ334-340 and sGC β1 Y135A R139A (24) were generated in the laboratory for this study. sGC β1 Δ265-271 was generated during a previous study in our laboratory (25). Human WT-IDO1-FLAG (Sino Biologicals # HG11650-CF), human WT-TDO-FLAG (Sino Biologicals # HG13215-CF), human WT-Hbβ-Myc-FLAG (Origene # RC203258) and human WT-Mb-Myc-FLAG (Origene # RC213272) mammalian expression plasmids were used. Heme binding studies in IDO1, Hbβ and Mb were performed using tetra cysteine tagged versions of the plasmids generated for this study. All the plasmids (WT and mutants) were confirmed by full plasmid sequencing before using in our experiments. Cultured cells were grown to 60% confluence before transfecting 10 cm dishes with 5 µg of each plasmid using Lipofectamine 2000 as per experimental groups. Protein expression was allowed for 48 h before harvesting cells and medium for further experiments.

### Immuno-precipitation (IP) and western blot

Cells were lysed using 50 mM HEPES pH 7.4 buffer with 100 mM NaCl, 0.1% Triton X-100, 5 mM Na-molybdate and EDTA-free protease inhibitor cocktail (Roche). Protein concentration was measured using the Bradford method (Bio-Rad # 500-0006). IP pull-downs were performed using 1 mg of cell lysate proteins with anti-FLAG antibody (Sigma # F1804), anti-sGC α antibody (Proteintech # 12605-1-AP), anti-sGC β antibody (Proteintech # 19011-1-AP) and anti-V5 antibody (Cell Signaling # 13202S). Each IP was performed using 4 µg of purified antibody. Protein G agarose beads (Millipore # 16-201) were used to pull down the antibody-protein complex. The beads were washed thrice with lysis buffer and SDS-PAGE sample buffer was added to the beads. For western blots to check protein expressions/IP’d and co-IP’d proteins across all samples in all experiments, we lysed cells as mentioned previously. In each input sample 50 μg of cell lysate proteins were boiled in Laemmli buffer, both input and IP samples were resolved onto 10% or 15% SDS-PAGE and transferred to PVDF membrane (Bio-Rad # 1620177) and probed for proteins of interest. Western blots were performed with anti-FLAG (Sigma # F1804; dilution 1:1000), anti-GAPDH (Proteintech # 60004-1-Ig; dilution 1:2000), anti-sGC α (Proteintech # 12605-1-AP; dilution 1:1000), anti-sGC β (Proteintech # 19011-1-AP; dilution 1:1000), anti-V5 (Cell Signaling # 13202S; dilution 1:1000), anti-Hsp90 (Proteintech # 13171-1-AP; dilution 1:1000), anti-IDO (Cell Signaling # 86630S; dilution 1:1000) and anti-α-Tubulin (Proteintech # 66031-1-Ig; dilution 1:2000). The proteins were detected using chemiluminescence using HRP conjugated secondary antibodies of either anti-mouse (Bio-Rad # 1706516, dilution 1:10,000) or anti-rabbit (Bio-Rad # 1706515, dilution 1:10,000) origin and ECL substrate (Thermo Scientific # 32106). The images were acquired using a chemidoc system from Bio-Rad.

### IDO1 and TDO activity assay from cell culture medium

The enzyme activity of IDO1 and TDO were measured using a colorimetric assay. The transfected cells were cultured with IDO1/TDO substrate L-Tryptophan at 2 mM in phenol red free DMEM medium containing 10% FBS for 12 h to allow for substrate utilization and Kyn product formation. The medium was collected and de-proteinized by adding an equal volume of 30% trichloroacetic acid (TCA) and incubated at 50 °C for 30 min. The tubes were centrifuged at 10,000 g for 10 min at room temperature to precipitate proteins from the medium. Equal volumes of de-proteinized sample were mixed with a freshly made 20 mg/ml solution of *p*-dimethyl-amino-benzaldehyde (Ehrlich’s reagent; Sigma # 109762) in glacial acetic acid at room temperature to allow for the formation of a yellow-colored product. The end point absorbance of this product was measured at 492 nm (Molecular Devices) to determine the concentration of Kyn in the medium. A standard curve was obtained using commercial Kyn (Sigma # K8625) dissolved in 0.5 N hydrochloric acid in various concentrations using the exact same method.

### Insertion of the tetra cysteine tag into human IDO1, Hb**β** and Mb genes

We used the commercially available mammalian expression plasmids for human IDO1, Hbβ and Mb mentioned previously as templates for inserting the TC tag which comprises “RWCCPGCCK” near the heme binding sites of each of these proteins. In IDO1 cDNA, A358 was deleted and in its position TC tag was inserted. In Hbβ cDNA, G84 was deleted and in its position TC tag was inserted and in Mb cDNA, G83 was deleted and in its position TC tag was inserted. All the TC tagged plasmid sequences were confirmed by whole plasmid sequencing.

### Staining live cells expressing TC-tagged heme proteins with FlAsH-EDT_2_ dye

COS-7 cells expressing TC-tagged heme proteins IDO1, Hbβ and Mb were grown in black walled 96-well plate (Greiner # 655090) in presence of 400 µM succinyl acetone (SA; Sigma # D1415) to heme deplete the cells for 2 days which were then stained with FlAsH-EDT_2_. Briefly, the medium was aspirated out from the wells and washed once with cell culture PBS. FlAsH-EDT_2_ (Toronto Research Chemicals # TRC-F335200) was dissolved in DMSO to a stock of 20 mM. The cells were stained with FlAsH-EDT_2_ diluted in opti-MEM medium at a final concentration of 5 μM at 37 °C for 30 min. The dye was aspirated out from the wells and washed twice with cell culture PBS. Phenol red free DMEM with or without D-ALA/Fe Citrate was added to the wells and the kinetic studies were performed.

### Live cell heme binding kinetics using FlAsH-EDT_2_ labeled heme proteins

The heme binding kinetics onto TC-IDO1/Hbβ/Mb under different experimental conditions was monitored in live cells stained with FlAsH-EDT_2_. We used Flex Station 3 (Molecular Devices) to perform kinetic studies over a 2 h period. We maintained a heme free control group in presence of SA to serve as a high baseline for each of the three TC-tagged heme proteins. The other groups had SA removed and D-ALA/Fe Citrate added to initiate endogenous heme production and subsequent heme insertion into the heme proteins over the 2 h period which would be indicated by quenching of their fluorescence signal upon heme binding. The FlAsH-EDT_2_ was excited at 508 nm and emission was recorded at 528 nm. The relative fluorescence units (RFU) readings were recorded from the bottom of the plate, at 100 reads per well per reading at medium sensitivity for the PMT settings for the entire duration of the experiment and the temperature was maintained at 37 °C inside the instrument.

### Proteasome inhibition studies using Bortezomib

Bortezomib (Cayman Chemicals # 10008822) was used to inhibit the proteasome in COS-7 and DU145 cells at a final concentration of 5 nM for 16 h during expression of IDO1, Hbβ and Mb proteins, and cell viability of both COS-7 and DU145 cells after undergoing the Bortezomib treatment was assessed based on their trypan blue exclusion.

## Statistical analyses

All experiments were done in three independent trials, with three replicates per trial. The results are presented as the mean of the three trial values ± standard deviation. The statistical test used to measure significance (*p*-values) was student’s t-test in the software Graph Pad Prism (v10).

## Acknowledgements

We thank Dr. Andreas Papapetropoulos (University of Athens, Athens, Greece) for providing the mammalian expression plasmids of rat sGC α1 and β1 and their mutants. We thank members of the Stuehr laboratory for the discussion of the manuscript.

## Data availability

All data are contained within this manuscript or are available from the authors upon request.

## Author contributions

P. B. and D. J. S. conceptualization; P. B., Y. D., D. T. J., and P. D. S validation; P. B., Y. D., D. T. J., P. D. S, S. M., and A. G. formal analysis; P. B., Y. D., D. T. J., and P. D. S investigation; D. J. S., S. M., and A. G. resources; P. B. and D. J. S. writing – original draft; D. J. S. writing–review & editing; P. B. and D. J. S. visualization; D. J. S. supervision; D. J. S. project administration; D. J. S. funding acquisition.

## Supporting information

This article contains supporting information.

