## Supplemental data for "Soluble guanylyl cyclase subunits act as Hsp90 co-chaperones to ensure the expression and functional maturation of hemeproteins in mammalian cells"

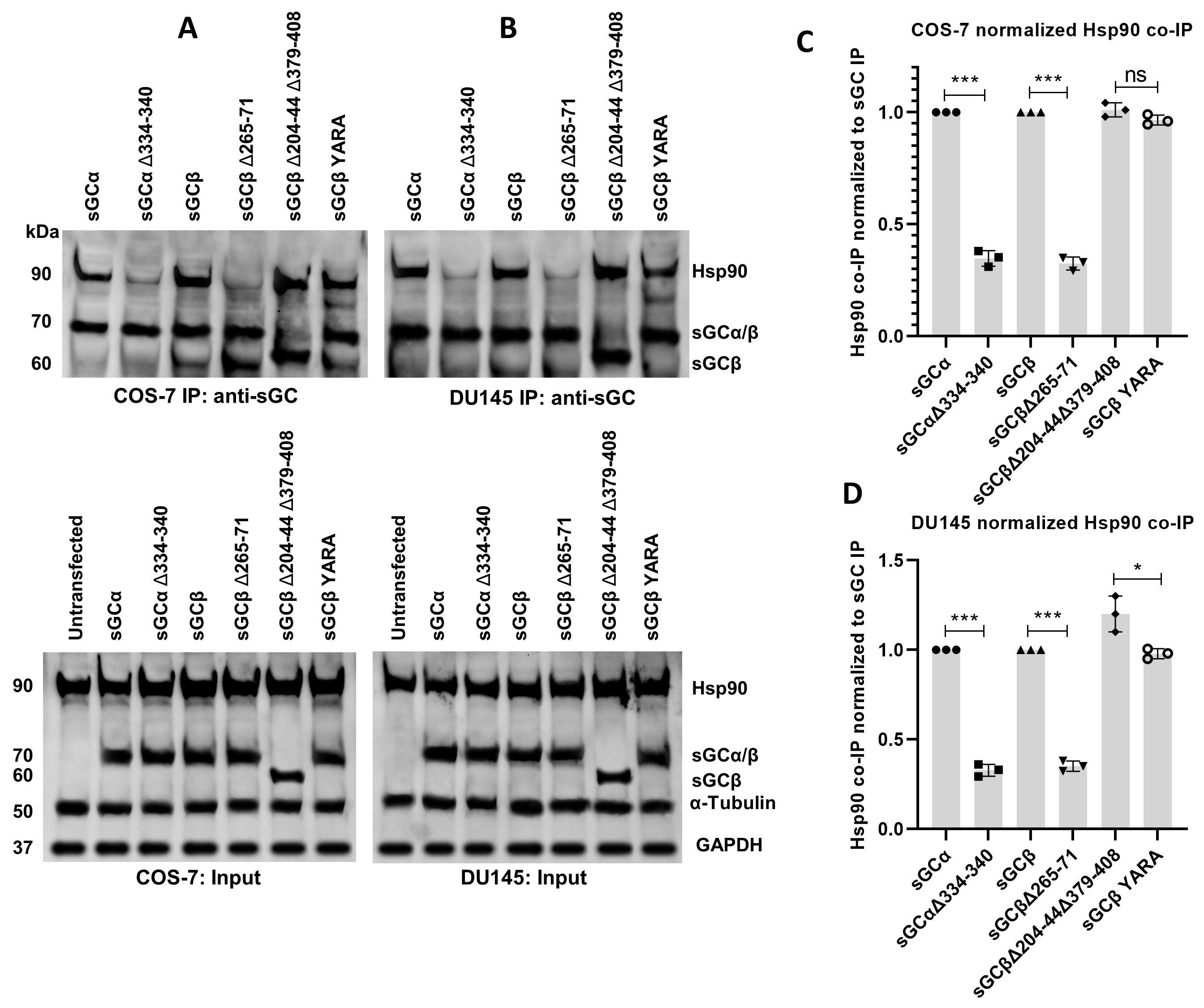

**Fig. S1: Interaction between sGC and Hsp90 in COS-7 and DU145 cells.** The indicated wild type and variants of sGCα and β subunits were expressed in COS-7 and DU145 cells. sGC was IP’d and the associated Hsp90 levels were compared across all the groups from the (A) COS-7 and (B) DU145 cells. The Hsp90-binding bands were analyzed by densitometry (C-D).

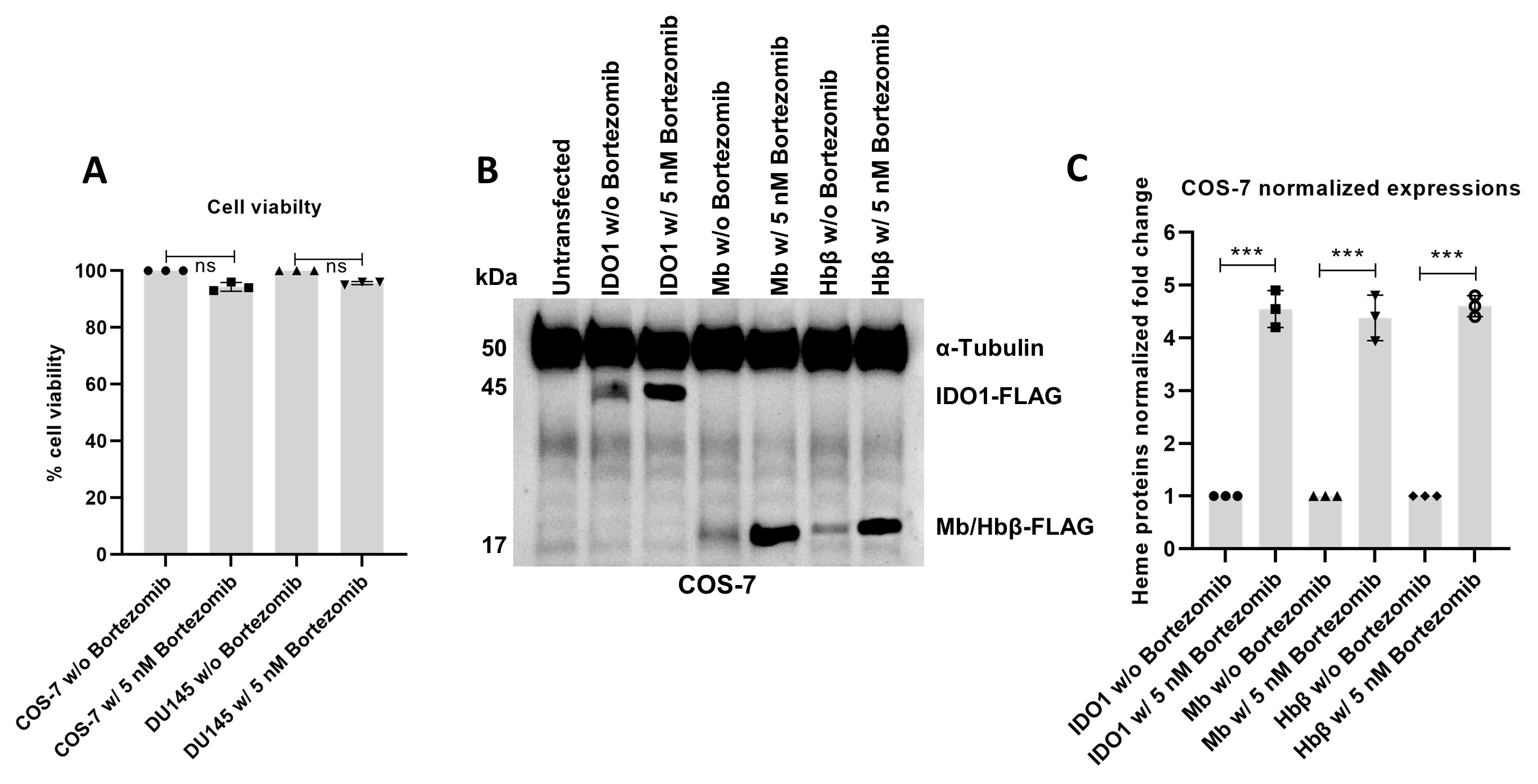

**Fig. S2: Bortezomib treatment for proteasome inhibition in COS-7 and DU145 cells.** The proteasome inhibitor Bortezomib was given to COS-7 and DU145 cells at 5 nM for 16 h to inhibit the proteasome while maintaining maximum cell viability (A). All heme proteins were found to accumulate inside the Bortezomib-treated COS-7 cells (B). The IDO1, Mb, and Hbβ expressions increased when compared to expression levels without Bortezomib treatment (C).
